# Evidence of ecosystem process recovery across a large-scale coral reef restoration programme using AI accelerated soundscape analysis

**DOI:** 10.1101/2025.09.24.678197

**Authors:** Ben Williams, Mars Coral Restoration Project Monitoring Team, Aya Naseem, Gaby Nava, Angus Roberts, Freda Nicholson, Aimee du Luart, Dave Erasmus, Ollie Stoole, Alistair Whittick, Rory Gibb, Tries B. Razak, Timothy A.C Lamont, Stephen D. Simpson, David J. Curnick, Kate E. Jones

## Abstract

1.

Coral reef restoration efforts are on the increase globally. However, reporting on the ecological outcomes of these efforts is rare and typically focuses on coral related metrics. As a result, understanding of whether restoration can recover broader aspects of reef functioning remains limited. In this study we use passive acoustic monitoring coupled with human-in-the-loop artificial intelligence to analyse >12 months of soundscape recordings from 45 sites across five biogeographically independent regions to investigate the impact of active restoration on reef functioning. We trained and rigorously evaluated machine learning models to identify 34 biological sound types within this data, generating >912,000 high-confidence detections. These detections were used to infer four key functions across healthy, degraded, early-stage (<3 months) and mid-stage (32–53 months) restored reefs. Restoration significantly enhanced: (i) biological sounds at night, key to recruiting juvenile fish; (ii) diversity of biological sounds, an indicator of fish community diversity; and (iii) snapping shrimp activity, an indicator of bioturbation. However, effects varied by region, and audible parrotfish grazing, key to algal control and bioerosion, did not differ among habitat types in four of the five regions. Our findings provide evidence that restoration can support recovery of broader ecosystem functioning when carefully implemented in the right contexts.

**Graphical abstract:** 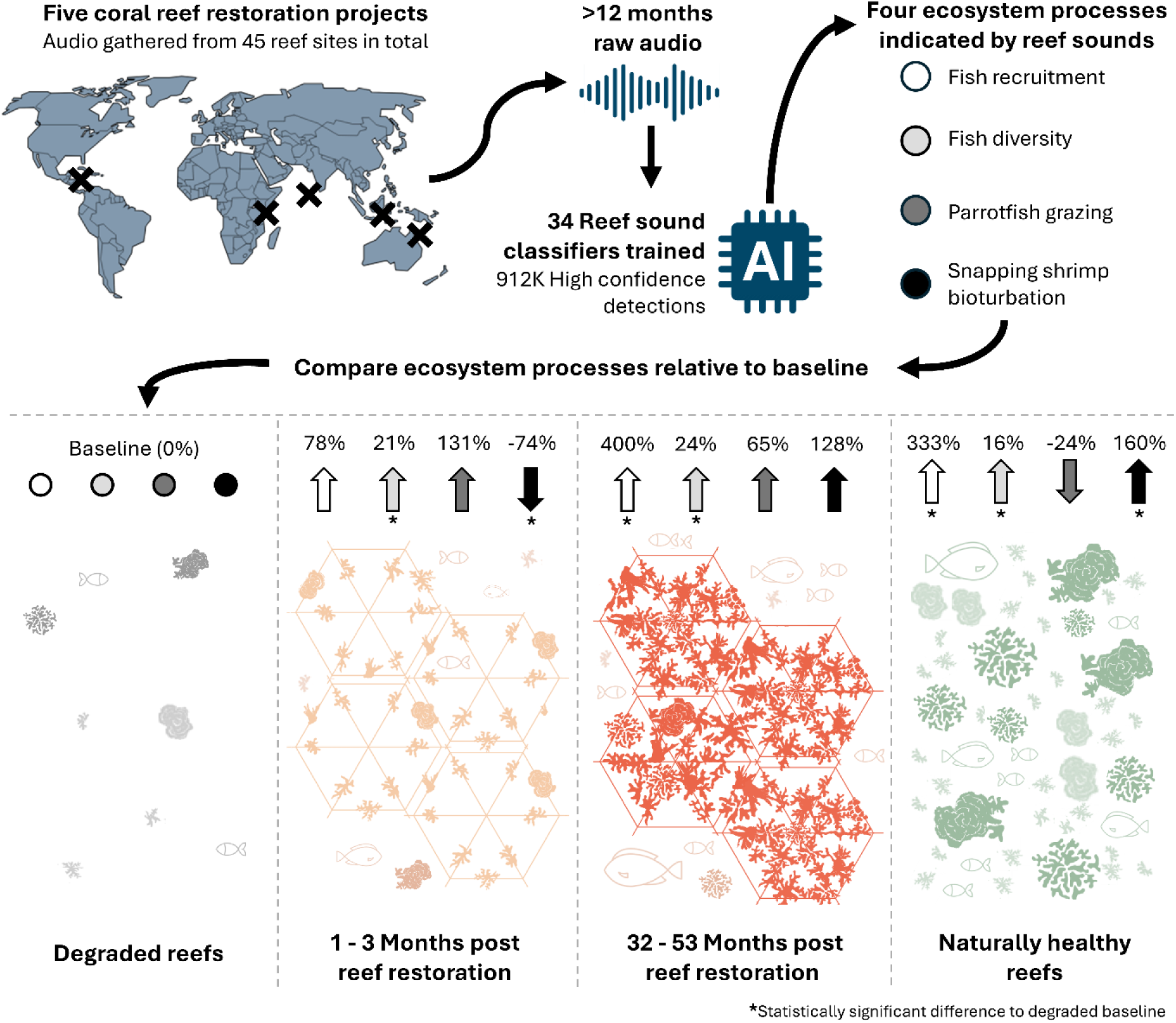

## 2. Introduction

Tropical coral reefs are among the most biodiverse ecosystems on Earth, supporting over 30% of known marine species and the livelihoods of around 500 million people^1,2^. Yet local and global anthropogenic stressors have reduced their capacity to deliver ecosystem services by more than 50% over the past 70 years^3^. In response, investment in reef restoration has expanded rapidly^4^. An English-language review documented 362 tropical reef restoration projects across the tropics as of 2019^5^, with a further 500 identified in Indonesia alone when sources in local languages were included^6^. Despite this surge, the ecological outcomes of reef restoration remain poorly understood. Furthermore, restoration is often implemented in areas where projects are highly likely to fail in response to local stressors or mid-century climate impacts^7^. Overcoming knowledge gaps in restoration outcomes is therefore essential for identifying its viability and the optimum strategies to achieve this across the full range of biogeographies and local contexts where coral reefs occur^8^.

Many restoration projects aim to restore a habitat’s ecosystem functioning towards pre-degradation conditions^9^. Ecosystem functioning here refers to the processes that mediate energy flow, material cycling, or community interactions, which collectively support an ecosystem’s services and resilience^10^. On coral reefs, functioning is underpinned by myriad ecological processes such as algal herbivory, habitat provisioning, larval recruitment and more^11^. Yet only a limited number of studies have examined the long-term recovery of ecosystem processes following coral reef restoration. For example, stabilisation and out-planting can restore carbonate budgets and small-scale complexity within 3-4 years, but often fail to recover large-scale structure^12–14^. Elsewhere, larval seeding has been shown to increase coral cover and fish abundance within three years^15^. However, such examples remain rare, meaning our understanding of whether restoration can re-establish broader ecological functioning and how this varies across approaches or regions remains limited^5^.

In this study, we focus on four key aspects of the reef soundscape that can serve as indicators of key ecosystem processes central to reef recovery: nocturnal fish sounds, biological sound diversity (phonic richness), parrotfish (*Scaridae*) grazing, and snapping shrimp (*Alpheidae*) activity. Each of these has well-established links to broader ecological processes. Nocturnal fish sounds represent the core component of the “recruitment cuescape”, the acoustic cues which attract the larval fish needed to replenish fish stocks^16–18^. Phonic richness quantifies the diversity of fish sounds present on a reef, which has been found to share a strong relationship with fish community diversity^19–21^, essential to broader reef functioning^22^. Grazing sounds from parrotfish are tied to algal herbivory and bioerosion, processes that maintain substrate for coral settlement and contribute to sand production^23–25^. Snapping shrimp are key agents of sediment bioturbation, providing an important pathway for nutrient cycling in otherwise nutrient-poor reef environments ^26,27^. Together, these acoustic indicators can be used to infer relative levels of critical ecosystem processes on reefs.

Monitoring reef ecosystem processes at scale is typically labour-intensive and costly. Consequently, monitoring of restoration outcomes is rare, and when undertaken, is usually limited to coral-specific metrics and seldom extends beyond 18 months^5^. Passive acoustic monitoring (PAM) with machine learning (ML) accelerated analysis offers a promising avenue to conduct efficient monitoring at scale^28^. Across a wide range of conservation settings in terrestrial systems, this technology is increasingly used to gather large ecoacoustic datasets with PAM recorders deployed in the field, which can then be rapidly analysed by ML algorithms^28,29^. However, this method remains comparatively underexplored on coral reef habitats, where applications of PAM are in their infancy and the use of ML to process these datasets has so far been limited to only a handful of studies^30–33^. Applying PAM and ML therefore presents a promising opportunity to advance understanding of ecosystem function recovery on coral reef restoration projects across the tropics.

Here, we test whether active coral reef restoration can drive the recovery of ecosystem processes beyond the coral community. We gather PAM data from five biogeographically isolated restoration projects, all applying the same widely adopted rubble stabilisation and coral out-planting methodology. On these data, we employ a cutting-edge human-in-the-loop machine learning pipeline to detect biological sounds. We use these detections to construct acoustic indicators of the fish recruitment cuescape, fish community diversity, parrotfish grazing, and sediment bioturbation processes. We compare these indicators across healthy, degraded, early-stage restored (<3 months) and mid-stage (32–53 months) restored reefs. We use this approach to test whether differences in ecosystem processes are observed between habitat types overall across the global dataset. Next, we take the same approach to report on outcomes of each individual restoration project in more granular detail. This allows us to test the hypothesis that healthy reefs exhibit higher levels of ecosystem processes than degraded reefs, and that restored reefs will show increases in these processes relative to degraded baselines.

## 3. Methods

### 3.1. Data collection

We gathered data from a total of 45 study sites across five reef restoration projects located in Australia, Indonesia, Kenya, Mexico and the Maldives (Fig. 1; Fig. S1). At each project, restoration was implemented following the Mars Assisted Reef Restoration System (MARRS) method^34^ (buildingcoral.com). This method targets former hard coral dominated habitat now characterised by unstable substrate, often comprised of coral rubble fields, which would otherwise require decades or longer to recover^35^. Hexagonal steel frames (“reef stars”) typically 0.91 m in diameter are coated in reef sand and fixed securely by divers to the substrate using stakes and cable ties. Fifteen coral fragments approximately 10 cm in size are attached to each of the reef stars prior to placement. A period of maintenance, typically comprising two to five visits separated by increasing intervals across the first 12 weeks post installation, is undertaken to remove algae and replace dead coral fragments, after which the system is left to recover without further intervention.

**Fig. 1:**
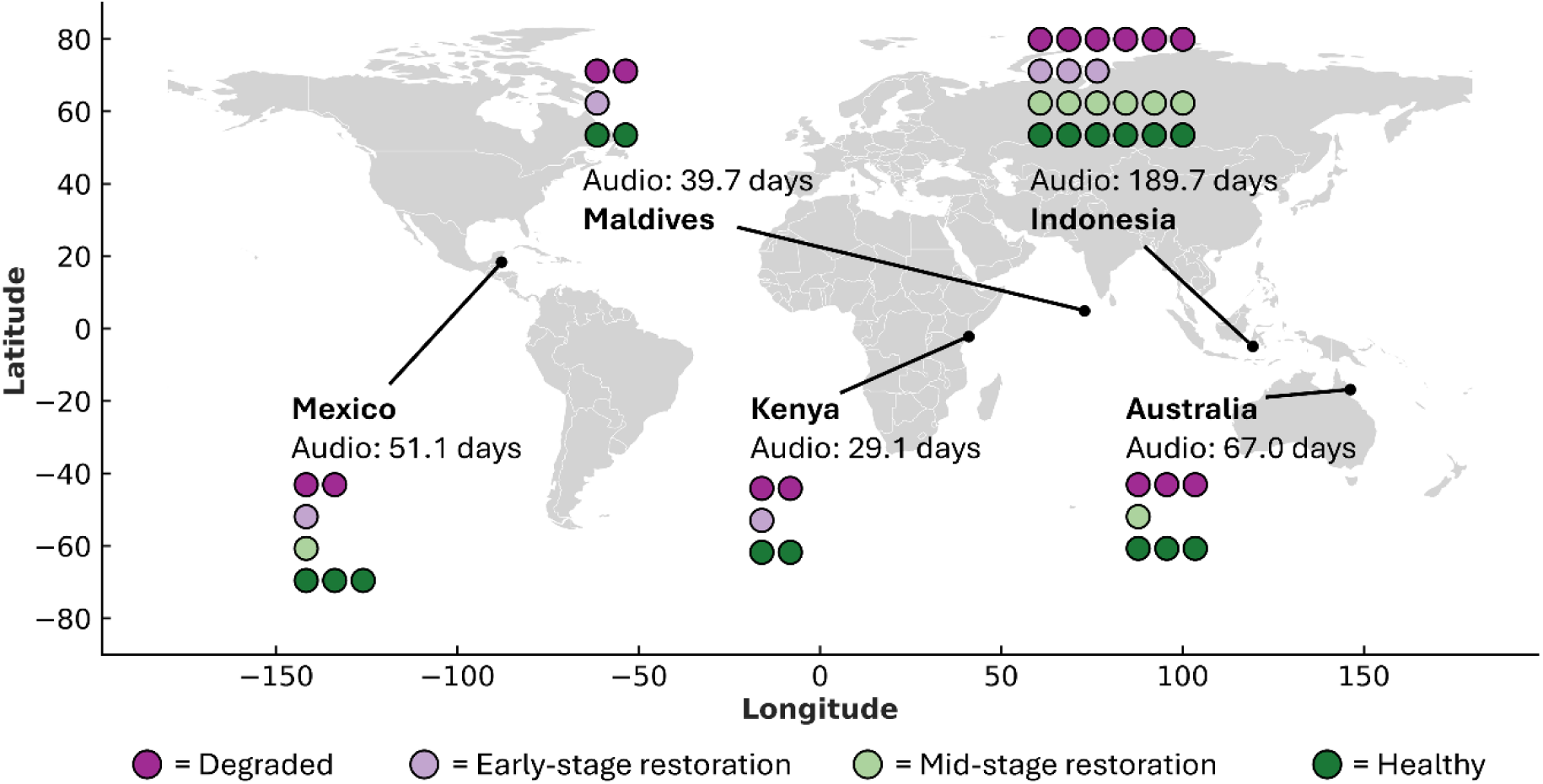
Map of restoration projects and count of habitat type replicates. Geographic locations of the five reef restoration projects used in this study. Coloured points indicate the number of study sites per habitat type within each project. Audio indicates the total duration of passive acoustic recordings taken across all sites within each project. A full map of each project is available in Supplementary 1.

Study sites were categorised into four habitat types: degraded, healthy, early-stage and mid-stage restoration. We defined the degraded category as reef habitat comparable to that of restored sites prior to the implementation of restoration. Healthy sites were defined as the least disturbed reef habitat in the local area; crucially this does not mean healthy controls were excluded from stressors entirely. We gathered basic ecological survey data from healthy and degraded sites to aid in their selection (Table S1). The early-stage restoration habitat type consisted of sites with 101-600 (65.7-390.5 m^2^) reef stars installed less than three months prior to acoustic data collection. The mid-stage restoration habitat type consisted of sites with 157-600 (102.3-390.5 m^2^) reef stars, installed 32-53 months prior to acoustic data collection. Given the full process of ecological succession and recovery is likely to take decades at a minimum on habitats such as coral reefs^35^, no plot was considered “late-stage” or “fully” restored. All sites within a project were selected to control for other variables, including reef zone, exposure and depth (which was between 1.2-4 m at low tide).All sites were comprised of at least 50 x 20 m of contiguous fringing reef habitat, and any two sites within each project were a minimum of 50 m apart.

We gathered acoustic data using HydroMoth audio recorders^36^ which were placed in the centre of each site and secured approximately 0.5m above the benthos using stakes. Care was taken to deploy devices at least 5 m from large barriers (e.g., boulders) which could cause occlusion of sounds. Each device was set to record at a sample rate of 16 kHz, using the lowest of the five gain settings with ‘low gain range’ enabled in the AudioMoth configuration application settings (v1.4). We set devices to record on a one in four-minute duty cycle, recording for a one-minute period followed by a three-minute break, except for the Indonesian project where recorders were set on a one in two-minute duty cycle with a battery change made during the study period. At each project, we gathered audio data simultaneously over an approximately one-month period. Any days where a device recorded <90% of its expected output, accounting for duty cycle, were discarded to remove incomplete days at the beginning and end of deployment. A mean of 27.85 (±10.11) unique days were recorded from each of the 45 sites across the full dataset (Table S1). All data collection was completed between October 2021 and June 2023.

### 3.2. Identifying sonotypes

Given the field of coral reef ecoacoustics is still in its infancy, the majority of biophonic sounds reported in studies to date cannot be attributed to specific taxa^20,37,38^. Instead, we categorised biophonic sounds into “sonotypes”, where groups of similar sounds that share consistent acoustic features (e.g., pitch, duration, pattern) are constructed^20,37,38^. We employed a conservative approach that restricted acoustically similar biophonic sounds into a single sonotype to mitigate “intergrading”, where variations of the same biophonic sound form a continuum rather than clear boundaries, which can undermine more granular groupings.

To construct sonotype groups, we identified unique sonotypes from each project individually using two approaches. The first approach (Fig. 2, 1c-6c) used unsupervised machine learning to screen the entire recording bank from each project. Here, all recordings were passed through an activity detector with a lenient threshold using the region of interest finder functionality in the scikit-maad package (v1.4.2) in Python (v3.10) (Fig. 2, 2c-6c). Parameters followed the packages default settings except for the minimum region of interest pixel count (min_roi) which was increased to 50 after trials revealed lower settings surfaced a high abundance of indistinguishable background sounds. The activity detector filtered out approximately 75% of each dataset. From the detected sounds, a two-second window was taken around each. These windows were then embedded using SurfPerch (Fig. 2, 3c), a pretrained neural network optimised to produce 1280-dimension embeddings of coral reef audio (Williams, 2025). To prepare each dataset for clustering, dimensionality reduction to 64 dimensions was first performed on the SurfPerch embeddings using Uniform Manifold Projection (UMAP)^39^ using a minimum distance (min_dist) of 0.1 and number of neighbours (n_neighbours) of 10 (Fig. 2, 4c). The resultant UMAP embeddings were then clustered by Gaussian Mixture Models (GMM) using the scikit-learn package (v1.5.1) (Fig. 2, 5c). The number of components (n_components) for the GMMs was set to 100, producing 100 clusters per dataset. Whilst this introduced redundancy by splitting some sonotypes into multiple clusters, it reduced the risk of acoustically distinct sonotypes being grouped together, which could result in rarer sonotypes being missed during subsequent manual inspection. From each cluster, 100 of the original two-second windows were then reviewed by an expert annotator (BW) to identify any prospective sonotypes present (Fig. 2, 6c).

**Fig. 2:**
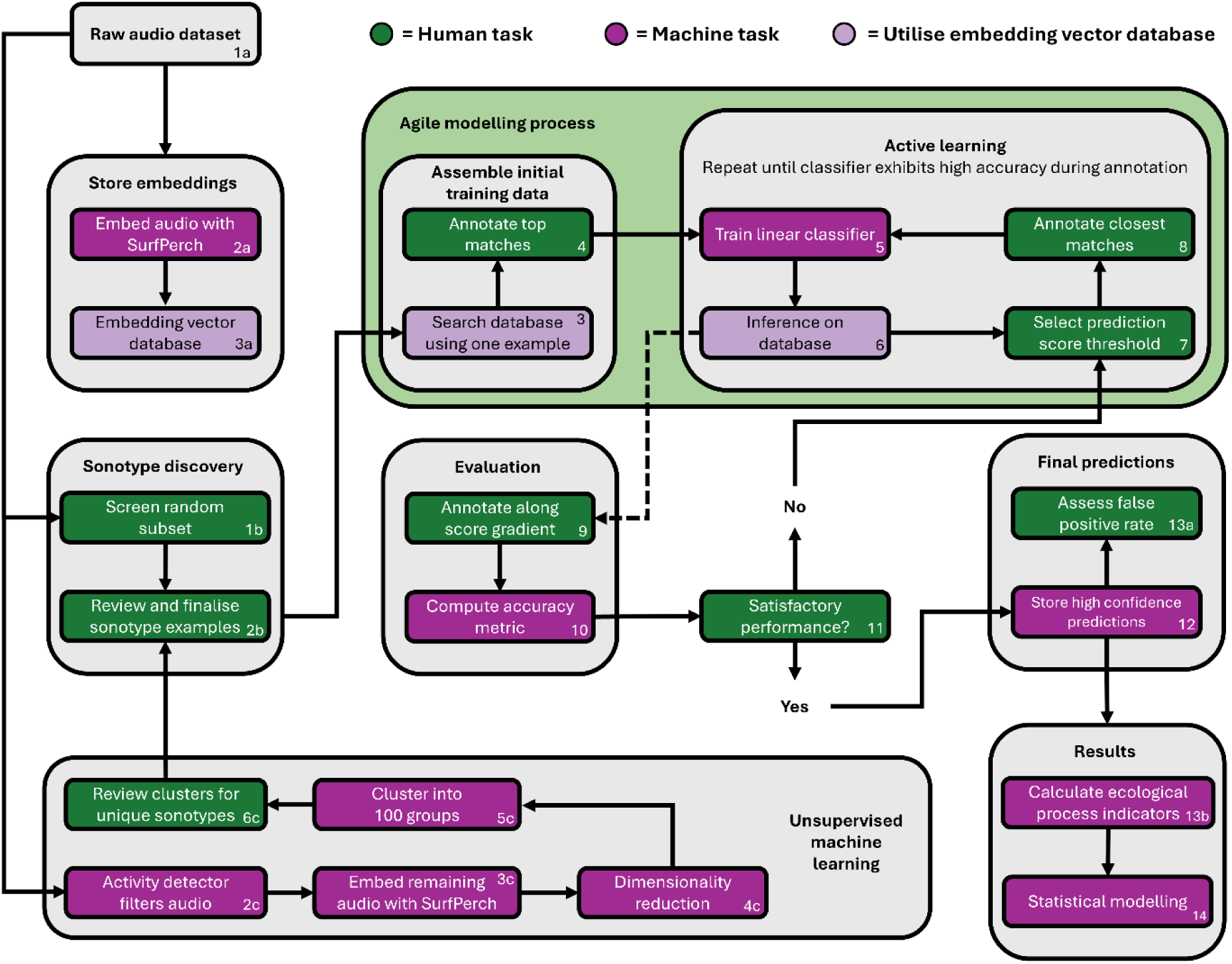
Analytical pipeline used to go from raw data to ecological process indicators. Flowchart summarising the analytical pipeline employed to identify sonotypes, generate detections of sonotypes with agile modelling, and model ecological functions from these detections. The agile modelling, evaluation and final prediction processes were completed separately for each sonotype within each project. Arrows indicate process flow; numbers denote the order of steps.

For the second approach for sonotype identification we performed a manual screening of 1000 one-minute files from each project taken at random, but balanced across sites, to check for any prospective sonotypes missed by the unsupervised learning (Fig. 2, 1b). These files were visually and acoustically screened by the human annotator to identify unique sonotypes. Across both the unsupervised and manual identification approaches, prospective sonotypes were scrutinised within and across the projects to group them into the final sonotypes used for further analysis (Fig. 2, 2b).

In total, we identified 15 unique sonotypes across all five projects (Fig. 3). The manual screening identified eight sonotypes, adding two to the 13 identified with the unsupervised ML approach. The array of sonotypes was broadly similar to those reported in existing work^20,37,38^. The Indonesian project exhibited the highest diversity of sonotypes, with ten sonotypes present, whereas Kenya and Mexico were tied for the lowest sonotype diversity at five. Only the “Snap” sonotype occurred across all five projects, with “Pulse knock” and “Scrape” found in four. Seven sonotypes were unique to an individual project. Notably “Scrape”, indicative of parrotfish grazing, was not found in the Kenyan dataset and hence this ecosystem process indicator could not be calculated for the Kenyan project.

**Fig. 3:**
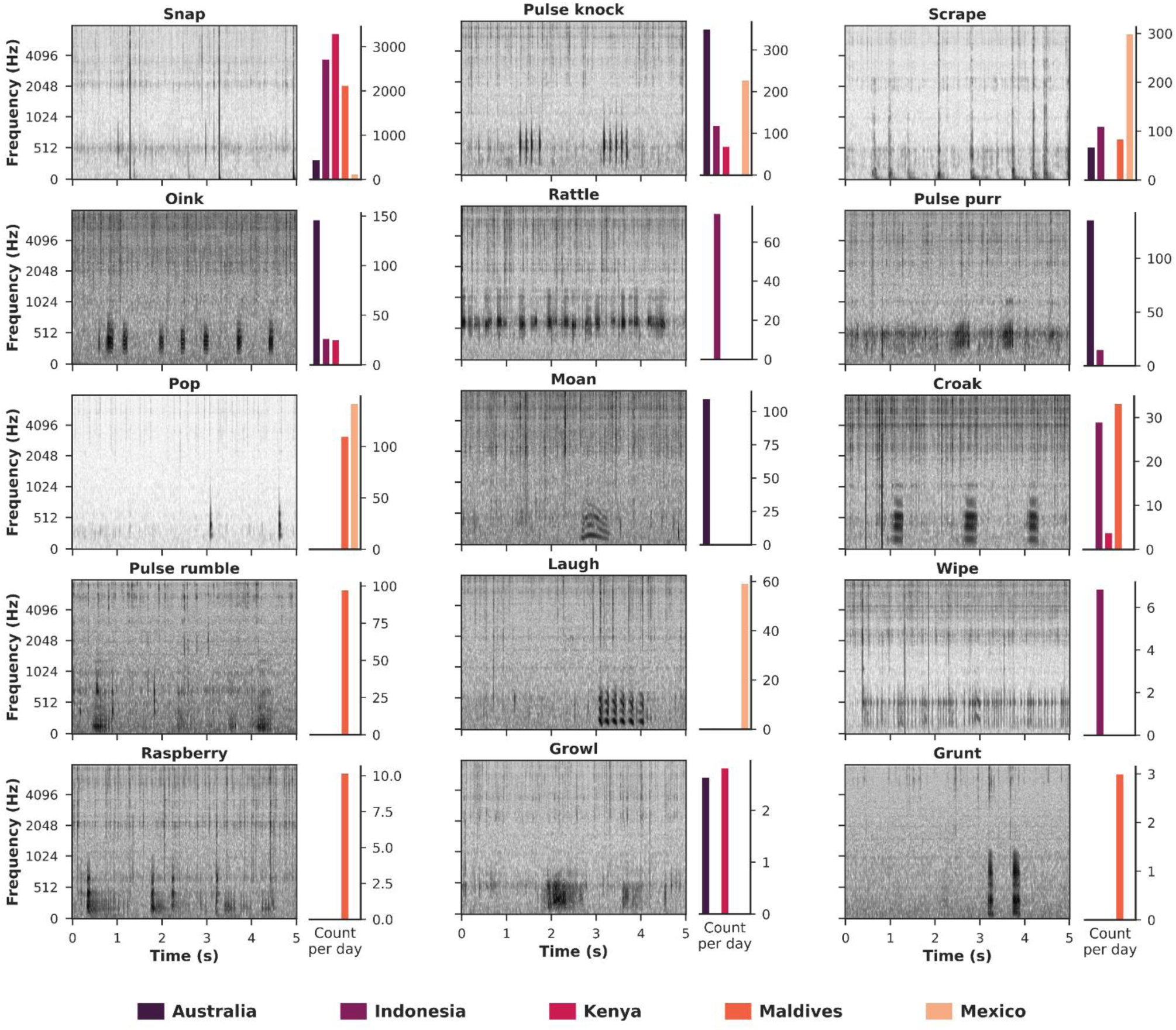
The fifteen sonotypes identified in this study. Log-mel spectrograms of audio samples containing the 15 sonotypes identified across the global dataset. Bar plots represent the mean number of high confidence detections (logit ≥1.0) per 24 hours of raw audio for each respective project. Sonotypes are sorted in descending order of total detection count across the full global dataset. Audio files and descriptions are available in Supplementary 1.

### 3.3. Generating sonotype detections with agile modelling

We followed an agile modelling approach to develop supervised machine learning classifiers for each sonotype, with models trained independently for each project to produce 34 country-specific classifiers (Fig. 2, 3-6). Agile modelling facilitates the rapid development of a classifier through leveraging human-in-the-loop annotation and model training, starting from a single sample of a given class^40,41^. Following Dumoulin et al., (2025), we started with a pretrained network, in our case SurfPerch, to embed the entire dataset of raw recordings from each project and store this in a vector database (Fig. 2, 2a). SurfPerch splits files into five-second windows, discarding any trailing audio where present, and produces a 1280-dimension embedding representation for each window^42^. Next, an example query of the target sonotype for the classifier is embedded and a nearest neighbour search against the vector database is performed using Euclidean distance (Fig. 2, 3). The top matches, typically 25–50 samples, were then labelled by a human annotator (BW) as a match or not, to the target sonotype (Fig. 2, 4). In rare cases where the human annotator could not determine if a sample was a match (e.g., due to low signal to noise ratio), the sample was left unlabelled.

The next stage of the agile modelling pipeline then involved an iterative active learning process (Fig. 2, 5-8). Here, I first trained a linear classifier on embeddings taken from each labelled sample, with a learning rate of 0.001 and batch size of 32 for 128 epochs (Fig. 2, 5). The trained model was then used to inference over the embedding vector database for the full dataset and a histogram of prediction scores (logit) produced (Fig. 2, 6). The 25-50 closest matches to a prediction score of 0 were first labelled (Fig. 2, 7-8), as these represent samples for which the model is most uncertain, and the classifier was then retrained with the addition of these samples. We repeated the model training and labelling process several times at a minimum for each sonotype during active learning (Fig. 2, 5-8). The prediction threshold used for labelling was shifted, using the histogram of scores as a reference, to surface more positive or negative matches for labelling as required.

Following the approach proposed in Navine et al., (2024)^43^, we then evaluated each resultant classifier one by one through inference across the embeddings of the full dataset, outputting a prediction score for each five-second window (Fig. 2, 6). The prediction scores were then ranked, and 25 samples were selected from each of four bins corresponding to the 0.9, 0.99, and 0.999 percentiles. These samples were then labelled blind by the human annotator (Fig. 2, 10). We calculated area under the receiver operator curve (AUC-ROC) from the prediction scores and annotations using bin-level weighting of inter-bin probability mass and direct within-bin score comparisons (Fig. 2, 11). Where AUC-ROC was low (typically <0.9), the agile modelling process was continued for at least three more iterations and evaluation performed again (Fig. 2, 11). Across all 34 classifiers trained using agile modelling, a mean AUC-ROC score of 0.975 (±0.029) was reported (Table S2). Despite high accuracy and recall on training and evaluation data, machine learning models can still exhibit low precision when deployed to detect rare events in large datasets—a common challenge in ecoacoustics, where the ratio of true positives to false positives is often low. To mitigate this, detections were filtered to only include high confidence detections, where the prediction score (logit) was ≥1 (Fig. 2, 12). This threshold provided a total of 912,131 detections across all 34 classifiers (Fig. 3; Table S2). To determine the rate of false positives that persist, for each classifier we performed a final evaluation on 100 randomly selected detections above the prediction threshold. The mean false positive rate after this evaluation was found to be 2.90% (±5.31) (Table S2).

### 3.4. Calculating acoustic indicators of ecosystem processes

The sonotype detections generated through agile modelling were used to calculate indicators for the relative levels of the four ecosystem processes identified. To quantify the fish recruitment cuescape, dusk-to-dawn periods were calculated using local sunset and sunrise times for each 24-hour period, via the Astral (v3.2) Python package. An additional 30 minutes was added before sunset and after sunrise to include twilight periods when fish recruitment may still occur. The nightly count of putative fish sonotype detections (which excludes snapping shrimp) was then multiplied by the recorder’s duty cycle to provide a score representing the abundance of nighttime fish sounds.

Phonic richness, a measure of sonotype diversity within the soundscape devised by Lamont et al., (2022), was used as an indicator for fish community diversity. Phonic richness was calculated as the number of unique sonotypes, excluding snapping shrimp, detected across all recordings at a site within a single 24-hour period (midnight to midnight).

Parrotfish grazing activity was estimated by counting detections of the “Scrape” sonotype within each 24-hour period at each site, and multiplying this by the recorder’s duty cycle. Snapping shrimp activity was quantified by counting detections of the “Snap” sonotype, within each 24-hour period at each site, and multiplying by the recorder’s duty cycle. To target nearby snaps, only occurrences with a high enough intensity to produce an unbroken fullband signal across 0-8 kHz were taken (Table S2), as opposed to distant snaps with breaks along this frequency range.

### 3.5. Statistical modelling

We used mixed models to test the relationship between each of the four ecological processes and habitat types. This modelling approach allowed us to test whether the relative levels of each ecological process, inferred from the soundscape, differed significantly between degraded, early-stage restored, mid-stage restored and healthy sites. This was performed across the whole dataset to identify the presence of any global trends, as well as for each project individually to report on specific outcomes from each project.

To reflect the hierarchical structure of the data, sites and their repeated daily recordings were nested within projects (countries). This specification allows correlation among observations from the same site across days and accounts for project-wide daily effects shared across sites (e.g., weather, lunar cycle). Random intercepts for project, site, and date were included to account for these sources of non-independence. Each function exhibited strong left skewness with overdispersion at high values. To account for this, we fitted a generalised linear mixed model (GLMM) with a negative binomial error distribution to each function individually. Each model included a fixed effect of habitat type and random intercepts for country, site, and date, with site and date both nested within project:

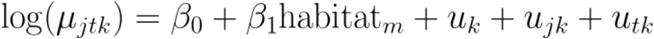

Here, μ_jtk_is the expected count for the ecological function (e.g., grazing rate) at site (*j*), on day (*t*), in country (*k*). β_0_represents the model intercept and β_1_*habitat* represents the fixed effect of habitat type (*m*). The u_*k*_ random effect accounts for variation among projects. The u_*jk*_ and u_*tk*_random effects account for variation among sites and dates respectively, each nested within project. For the fish recruitment cuescape, an additional term was added to account for the variability in length of nighttime periods across locations between projects and dates by passing the log of the maximum possible score for each night (*u*_*tk*_^*max*^) as an offset to the count. For project-specific models, the same structure was used with the project-level random effect excluded; only site and date were included as random effects without nesting. All models were implemented in R using the lme4 package (v1.35.5).

## 4. Results

### 4.1. Global responses of ecosystem process indicators across habitat types

Across the global dataset, we found a significant difference between degraded reefs and one or more of the other habitat types (early-stage restoration, mid-stage restoration, healthy) for the fish recruitment cuescape, phonic richness and snapping shrimp activity (Fig. 4). The most dramatic increase compared to degraded reefs was observed for the fish recruitment cuescape, with a 400% increase (β = 1.61, *p* < 0.001) on mid-stage restored reefs and a 333% higher value (β = 1.47, *p* < 0.001) on healthy reefs; the healthy and mid-stage restored reefs did not differ significantly from one another in post-hoc analysis (β = 0.144, *p* = 0.75), indicating similar overall values. We found early-stage restored reefs trended towards an increase relative to degraded reefs, but this was non-significant. Phonic richness exhibited a smaller, yet comparatively consistent, increase over degraded reefs for the three other habitat types, with a 16% (β = 0.15, *p* = 0.007) higher value on healthy reefs, alongside a 24% (β = 0.22, *p* = 0.009) and 21% (β = 0.19, *p* < 0.004) increase on early-stage and mid-stage restored reefs respectively. Finally, we found snapping shrimp activity was significantly lower on early-stage restored reefs compared to degraded reefs, with a 74% decrease (β = - 1.29, *p* = 0.025), whereas it was significantly higher on healthy reefs by 160% (β = 0.96, *p* = 0.024), with a non-significant increase reported for mid-stage restored reefs. The only acoustic indicator which did not exhibit differences across habitat types was parrotfish grazing, where non-significant trends towards an increase on the early and mid-stage restored reefs, and a decrease on the healthy reefs, were observed relative to the degraded reefs.

**Fig. 4:**
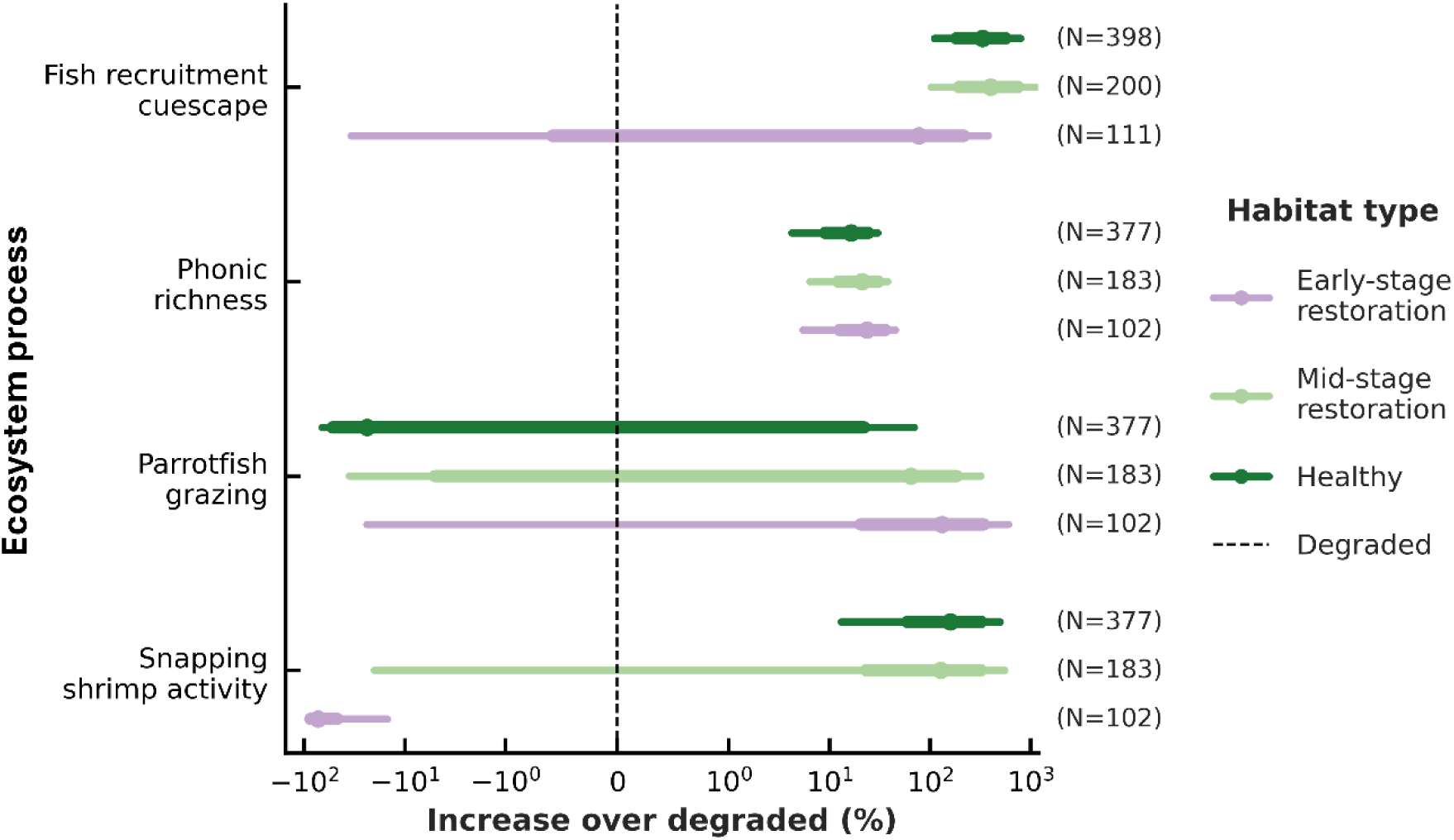
Impact of restoration on ecosystem process indicators across all projects. Percentage change in acoustic indicators of ecosystem processes for habitat types against the degraded baseline (log-scale), reported by the GLMM. Points indicate mean estimated increase, thick whiskers represent 75% confidence intervals, thin whiskers represent 95% confidence intervals. N values represent the number of samples (unique days) within each respective habitat type for the given ecosystem process.

### 4.2. Project-specific responses of inferred functions across habitat types

In the Australian project, we found the mid-stage restored reef showed a 91% increase in the fish recruitment cuescape relative to degraded reefs (β = 0.65, *p* = 0.002), whereas the healthy treatment exhibited only a non-significant 29% increase (β = 0.26, *p* = 0.08) (Fig. 5). For phonic richness, healthy reefs exhibited an 18% reduction compared to degraded reefs (β = - 0.20, *p* < 0.001), with no significant change on the mid-stage restored reefs. Australia was also the only project for which we observed any significant difference in parrotfish grazing, where this was found to be lower on healthy reefs and the mid-stage restored reefs which reported 73% (β = - 1.33, *p* < 0.001) and a 62% (β = - 0.97, *p* = 0.045) lower values respectively relative to degraded reefs.

**Fig. 5:**
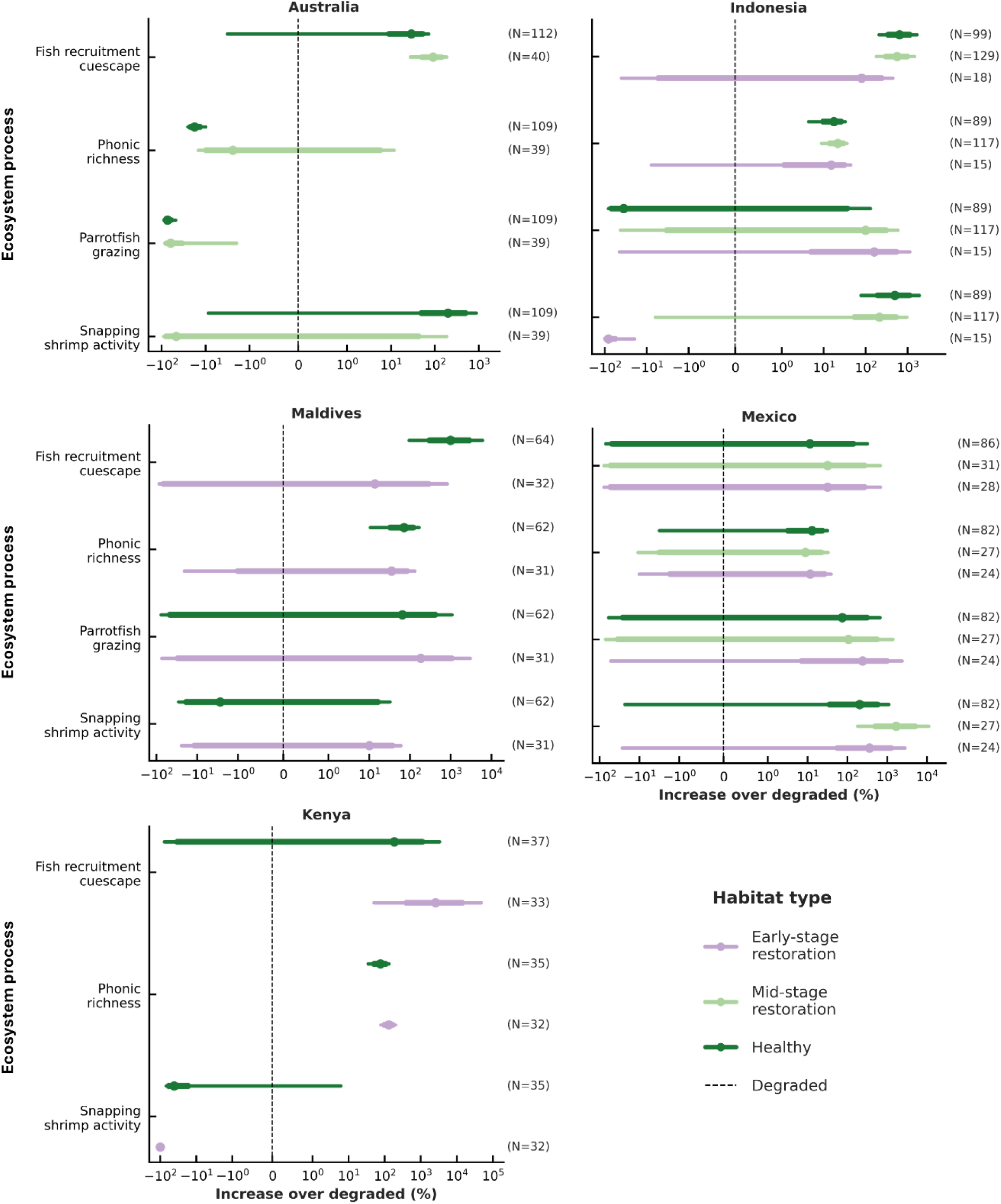
Local impacts of restoration on ecosystem process indicators. Percentage change in acoustic indicators of ecosystem processes for habitat types against the degraded baseline, reported by comparisons with the GLMM for each project individually. Points indicate the estimated increase, thick lines represent 75% confidence intervals, thin lines represent 95% confidence intervals. N values represent the number of samples (unique days) within each respective habitat type for the given ecosystem process.

In the Indonesian project, we found both the healthy and mid-stage restored reefs had markedly enhanced fish recruitment cuescapes relative to degraded sites, with increases of 639% (β = 2.00, *p* < 0.001) and 564% (β = 1.89, *p* < 0.001) respectively. Healthy and mid-stage restored treatments also had significantly increased phonic richness scores, 18% (β = 0.16, *p* = 0.008) and 22% (β = 0.20, *p* = 0.001) higher than degraded reefs respectively. In contrast, we found snapping shrimp activity was significantly higher on healthy reefs, with a 496% increase (β = 1.79, *p* = 0.004) over degraded reefs, and mid-stage restored reefs showed a near significant 214% increase (β = 1.15, *p* = 0.063), but snapping shrimp activity was significantly lower (- 82%; β = - 1.70, *p* = 0.024) on early-stage restored reefs compared with degraded reefs.

The projects in the Maldives and Mexico showed less definitive differences. In the Maldives project, we found only healthy reefs exhibited significant differences from the degraded reefs, with a nearly tenfold increase in the fish recruitment cuescape (993%; β = 2.39, p = 0.006) and a 72% higher phonic richness score (β = 0.54, p = 0.016) relative to degraded reefs. The early-stage restored reef showed positive trends across all acoustic indicators, but none reached significance, and no mid-stage restored reefs were available to test. In the Mexico project, whilst we observed healthy, early-stage and mid-stage restored reefs exhibited trends towards an increase in all four acoustic indicators, none of these were significant except for snapping shrimp activity on the healthy reefs, which was 1669% higher (β = 2.87, p = 0.0022) than the degraded reefs.

In the Kenyan project we observed the fish recruitment cuescape on the early-stage restoration reef was dramatically increased by about 2573% compared to degraded sites (β = 3.29, *p* = 0.025), though no significant difference was observed between healthy and degraded reefs. Phonic richness was 77% (β = 0.57, *p* < 0.001) and 131% (β = 0.84, *p* < 0.001) higher on healthy and early-stage restoration reefs. We found snapping shrimp activity exhibited a large reduction on the early-stage restoration site with a decrease of 99% (β = -4.33, *p* < 0.001) compared to degraded sites. We did not observe any parrotfish grazing, characterised by the “Scrape” sonotype, across any habitat type in the Kenyan dataset (Fig. 3).

## 5. Discussion

Our results indicate that active coral reef restoration can initiate the recovery of broader ecosystem processes which underpin reef functioning. Specifically, interventions that implement substrate stabilisation combined with coral out-planting at scale can lead to the recovery of processes beyond the rubble and coral assemblage these target, without further intervention. However, we find responses to restoration vary considerably across the five biogeographically isolated projects tested. We also find that indicators of these processes do not always differ significantly between degraded reefs, representative of the reefs targeted for restoration, and the healthiest baselines available locally.

Looking to individual processes, we observe evidence that active restoration can rebuild a diverse fish community essential to reef functioning. We found the fish recruitment cuescape was severely impacted on degraded reefs when compared with healthy baselines. However, by the time reefs reach mid-stage restoration (32–53 months) the recruitment cuescape can recover to levels comparable to that of healthy baselines. Furthermore, phonic richness was significantly reduced on degraded reefs when compared to healthy reefs, but recovery to levels comparable to that of healthy baselines on both early-stage (< 3 months) and mid-stage restored reefs was observed. Together, these two indicators provide evidence that active restoration can re-establish the acoustic cues needed for larval settlement and the subsequent return of a diverse fish community.

In addition, we find evidence that active restoration can rebuild snapping shrimp activity which contribute to nutrient cycling through sediment bioturbation. This indicator exhibited an initial reduction against the degraded baseline for early-stage restoration plots, followed by partial recovery on mid-stage plots. This initial reduction may be explained by the disturbance to the substrate post-installation of reefs stars which stabilise and cover the benthic habitat inhabited by snapping shrimp. However, snapping shrimp activity varied among projects, with degraded reefs in the Maldives and Kenya exhibiting higher, though non-significant, levels. Parrotfish grazing was the only function which did not report a meaningful difference across any habitat type. Possible explanations include individual parrotfish graze across spatial scales larger than the sites sampled in this study (50 x 20 m of contiguous fringing reef habitat type), compensatory increases in grazing intensity where abundance is lower, or because grazing on some substrates (e.g., rubble versus live coral) is less audible.

High variability in the inferred levels of ecosystem processes between the four habitat types across projects could be due to a multitude of factors. Firstly, each project is in a different biogeographic realm, with unique ecological assemblages that may respond differently to the same intervention. These projects were subject to a range of initial causes of reef degradation and exhibited differing upper limits of reef health within the region (Table S1). For example, the Indonesian project is situated within the Tropical Indo–West Pacific, the most biodiverse realm of coral reefs globally^44^, and was subject to locally acute degradation caused by bomb fishing. The mean coral cover on healthy and degraded controls exhibited stark differences at 71.8% (± 10.1%) and 10.2% (± 8.0%) respectively. In contrast, the Mexican project is situated within the Tropical West Atlantic realm, estimated to have under half the biodiversity of the Tropical Indo–West Pacific^44^. The Mexican project was subject to more geographically pervasive degradation through a combination of historical overfishing, coral disease and bleaching (Table S1). Here, mean coral cover of healthy and degraded reefs was less contrasting at 20% (± 7.9%) and <5% respectively, indicating a lower quality of maximally healthy reefs in the area. The only acoustic indicator that differed between healthy and degraded reefs in the Mexican project was phonic richness, with no recovery for this observed on the early or mid-stage restored reefs.

This study strongly supports the importance for reef managers of considering how strongly degraded sites targeted for restoration diverge from locally healthy baselines^7^. Where significant differences exist, we found evidence to suggest active restoration can successfully initiate or even lead to full recovery of some ecosystem processes within relatively short (<5 year) timeframes. Where differences are not observed, it is important to consider the reasons behind this. Lack of observable differences may occur because degradation events have not been severe enough to reduce reef health to levels where restoration is required. Alternatively, degradation may be so pervasive that ecological homogenisation has occurred, with high-quality baselines no longer present. For example, phase shifts to algal-dominated states, driven by eutrophication or the loss of habitat following overexploitation, can create reinforcing traps that prevent natural recovery^45^. In such systems, active restoration alone is unlikely to succeed, and broader interventions such as fisheries management or pollution control may be required. Additionally, climate change now represents the dominant long-term threat to coral reef systems^46^. Reef managers should therefore consider the predicted impacts of climate related stressors present on their reefs when deciding whether active restoration offers a viable strategy^7^, which could be estimated through resilience surveys prior to implementing restoration^47^.

The use of passive acoustic monitoring (PAM) to assess potential gains in ecological functions on restored reefs shows promise, but there remain further opportunities to develop this approach. Growing efforts to link biophonic sounds to specific taxa or ecological processes will likely help unlock the ability to infer additional functions using PAM^48,49^, such as levels of predation, or seasonally variable processes including courtship and reproduction, and formation of spawning aggregations^23,50^. Assessing additional functions could help provide a more thorough understanding as to whether active restoration leads towards the recovery of self-sustaining ecological systems comparable to healthy reefs in the local area, or whether some key functions fail to return, or diverge from naturally healthy baselines without additional interventions. Long-term temporal sampling on the scale of years could be used to better understand the timeframe over which recovery occurs and how this differs across inferred functions through early to late-stage recovery. The method presented in this study for assessing functioning through ecoacoustics could be applied to assessing the efficacy of alternative restoration strategies, as well as other conservation interventions (e.g., wastewater management, fishing restrictions, etc.) and measures to limit degradation pressures (e.g., pollution, storm events, etc).

## 6. Conclusion

We found that active restoration can lead to the onset or full recovery of important ecological processes on tropical coral reefs. Interventions combining coral out-planting with rubble stabilisation can generate ecological benefits that extend beyond these two components alone, supporting the recovery of the broader ecosystem functioning needed for a self-sustaining reef assemblage. However, careful consideration is needed when deciding if active restoration is the best approach. Local context, including the causes and extent of degradation as well as projected future impacts and reef resilience, will ultimately determine whether healthy reefs can persist in a given area. We also demonstrate the utility of AI with human-in-the-loop led analysis to produce ecological insights from large ecoacoustic datasets, with relevance to ecologists working across a broad range of habitats.

## 7. Data availability

The full set of raw recordings taken throughout the study, audio files of all detections output by the agile modelling process, and an explanation on how to use this data are all available in a Figshare repository: https://doi.org/10.5522/04/29958062.

## 8. Code availability

All code used to complete the study is available in a GitHub repository: https://github.com/BenUCL/MARRS_global_acoustic_study. To implement the agile modelling process used, we recommend the Perch Hoplite Repository: https://github.com/google-research/perch-hoplite.

## 9. Acknowledgements

Funding was provided by a Fisheries Society of the British Isles PhD Studentship (awarded to B.W); a research fellowship from the 1851 Royal Commission (awarded to T.A.C.L.); an international travelling fellowship from the Fisheries Society of the British Isles (awarded to T.A.C.L. and T.B.R); and a Pew Fellowship in Marine Conservation (awarded to T.B.R).

In gathering the Australian dataset, we thank Mars Sustainable Solutions for supporting restoration and data collection. In Australia, we wish to acknowledge and thank the Gunggandji people as the Traditional Custodians of the Sea Country where our research took place. We also thank Reef Magic and GBR Biology for permissions to implement restoration and gather data under the permits: G20/42902.1

For the data gathered from Kenya, we wish to thank the Ocean Trust for leading the fieldwork and access to their restored system, as well as Lamu County for granting permissions to gather this observational data as part of the Ocean Trust monitoring effort.

Data in Indonesia were collected as part of the monitoring program for the Mars Coral Reef Restoration Project, in collaboration with Universitas Hasanuddin. We thank Lily Damayanti, Pippa Mansell, and Mars Sustainable Solutions team for support with fieldwork logistics and monitoring efforts. Logistical research support was provided by Mars Sustainable Solutions. We also thank the Department of Marine Affairs and Fisheries of the Province of South Sulawesi, the Government Offices of the Kabupaten of Pangkep, the community of Pulau Bontosua for their support. Fieldwork in Indonesia was conducted under an Indonesian national research permit issued by BRIN (number 108/SIP/IV/FR/2/2023, jointly held by T.A.C.L. as lead foreign researcher and T.B.R. as lead Indonesian host researcher, with all foreign researchers named on the permit), with ethical approval from BRIN and Lancaster University. We thank Prof J. Jompa and Prof R.A. Rappe at Universitas Hasanuddin for logistical assistance with permit and visa applications.

For the data gathered from the Maldives we wish to thank staff of the Maldives Coral Institute implementing the restoration plots and gathering data, as well as the Maldives Environmental Protecton Agency for providing a decision statement approving the restoration project and gathering of monitoring data under (203-ECA/PRIV/2021/337).

For the data gathered from Mexico, we wish to thank the staff of Oceanus A.C. for implementing the restoration plots and gathering data, as well as the National Commission of Protected Areas (CONANP) who administer the protected area through the National Park Reefs of Xcalak, and the help of the Xcalak Local Restoration Group for helping on the restoration activities. Restoration and the gathering of monitoring data permissions were granted by the General Directorate of Wildlife, under the Undersecretariat of Environmental Policy and Natural Resources permissions under permit (23/K5-0064/07/23).

## 11. Supplementary 1

### Study sites

Basic maps of study sites made with the Python’s Contextily package (v1.6.2). can be found below. However, we recommend readers access the Google Earth project available online where an interactive map of each project and labelled sites can be explored by navigating the table of contents at: https://tinyurl.com/3xsvf5jt. Alternatively, the *study_sites_map.kml* file in the supplementary repository can be used to reproduce the project in other common mapping software’s such as QGIS, CesiumJS or Garmin BaseCamp.

### Australia

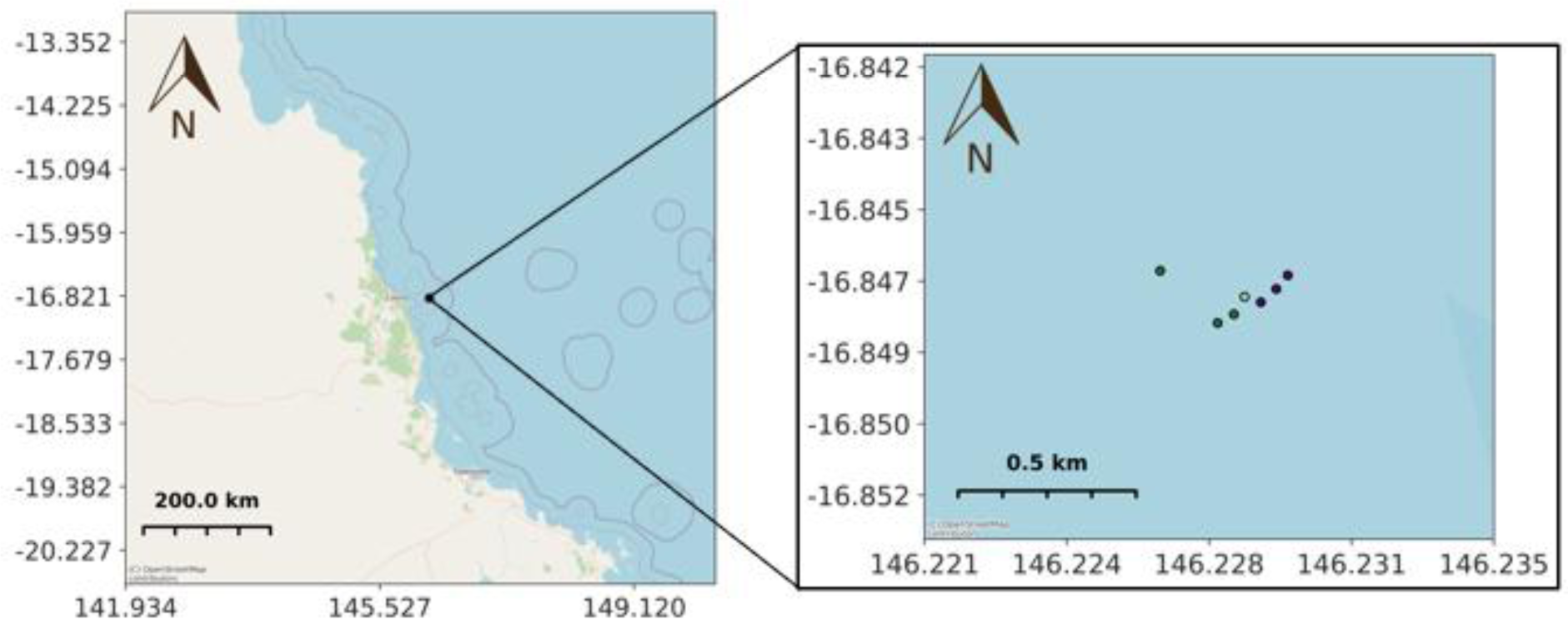

### Indonesia

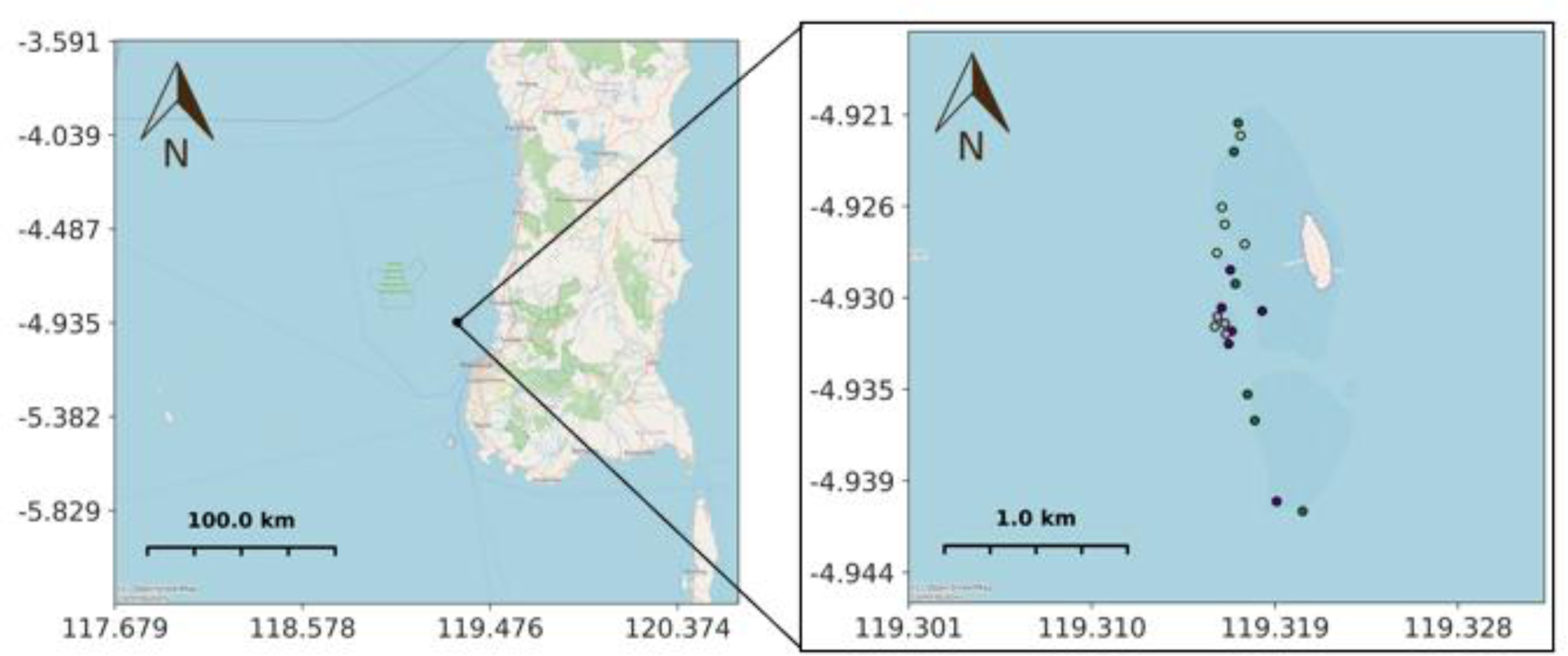

### Kenya

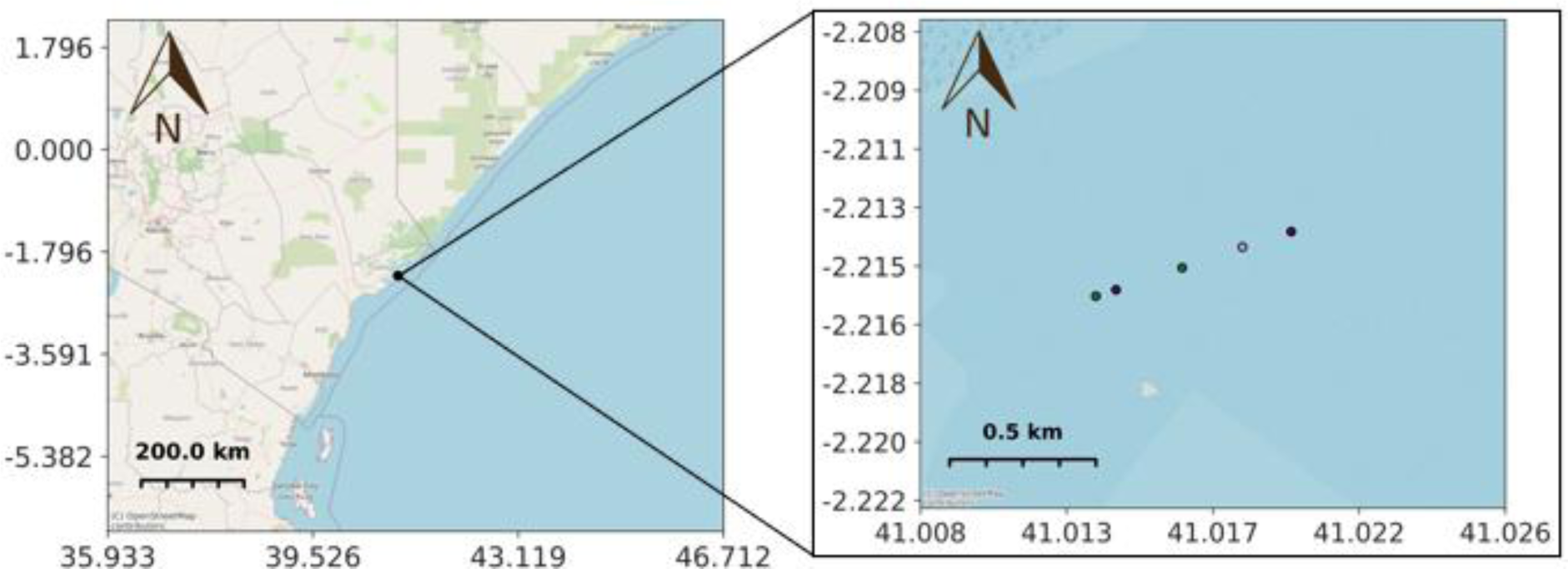

### Maldives

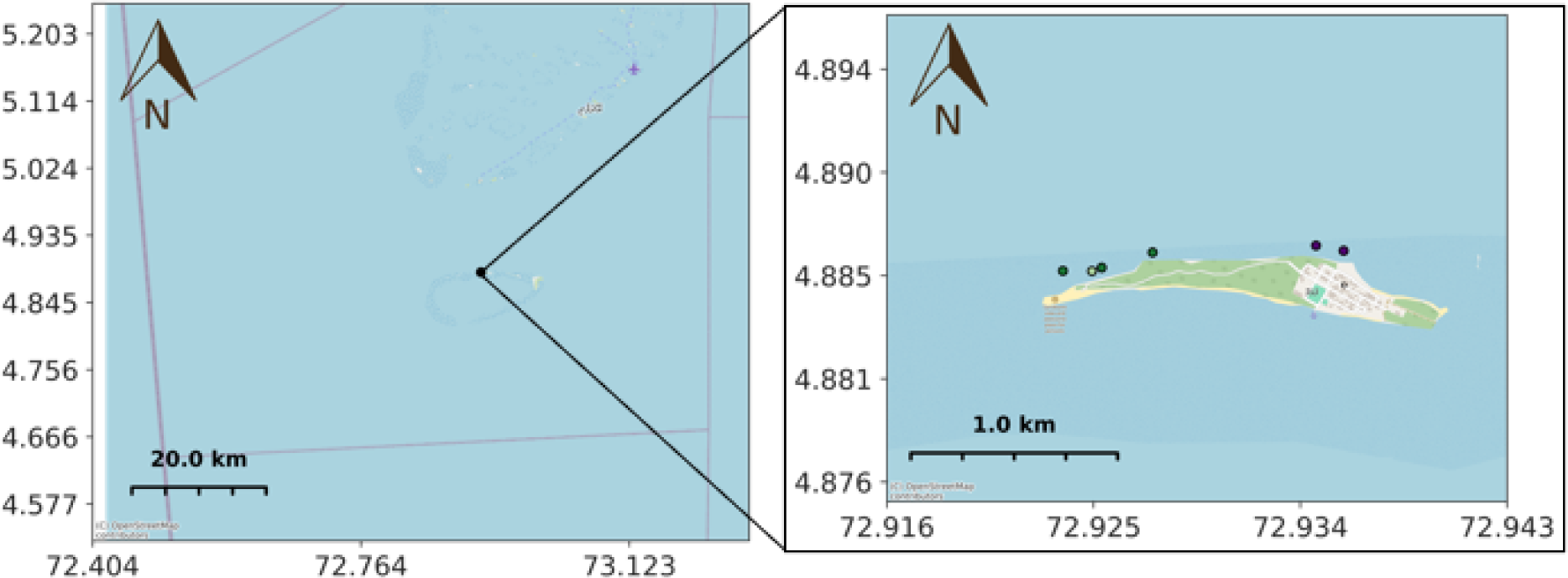

### Mexico

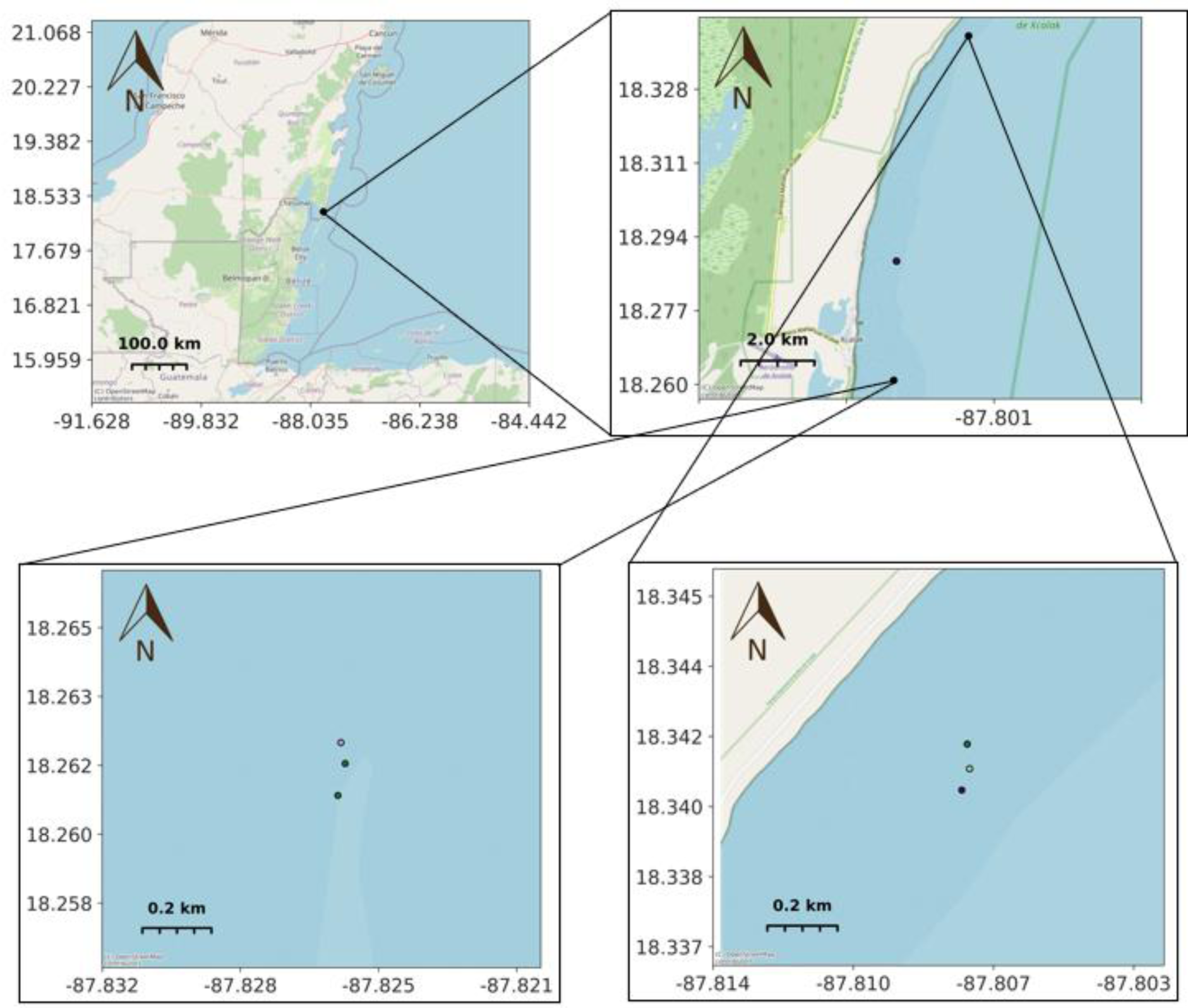

**Supplementary Table 1:** Metadata from each site from which acoustic data were gathered. Age of restoration plot indicates the number of months prior to acoustic data collection that the restoration plot was implemented. Cause of degradation indicates the primary drivers of reef degradation on the sites within each project. Supporting survey data included visual census data gathered by each project to support their selection of healthy and degraded baselines. Days recorded indicates the total number of unique calendar days each recorder was operating across during its duty cycle.

| Project | Site | Habitat type | Number of reef stars on plot | Age of restoration plot | Cause of degradation | Supporting survey data | Days recorded |
| --- | --- | --- | --- | --- | --- | --- | --- |
| Australia | D1 | Degraded | NA | NA | Cyclone damage | Hard coral cover = 25% | 39 |
| Australia | D2 | Degraded | NA | NA | Cyclone damage | Hard coral cover = 21% | 40 |
| Australia | D3 | Degraded | NA | NA | Cyclone damage | Hard coral cover = 8% | 40 |
| Australia | H1 | Healthy | NA | NA | NA | Hard coral cover = 77% | 39 |
| Australia | H2 | Healthy | NA | NA | NA | Hard coral cover = 43% | 36 |
| Australia | H3 | Healthy | NA | NA | NA | Hard coral cover = 65% | 40 |
| Australia | R1 | Mid-stage restored | 439 | 32 months | Cyclone damage | Hard coral cover = 55% | 41 |
| Indonesia | D1 | Degraded | NA | NA | Bomb fishing, coral mining | Hard coral cover = 12% | 26 |
| Indonesia | D2 | Degraded | NA | NA | Bomb fishing, coral mining | Hard coral cover = 5% | 25 |
| Indonesia | D3 | Degraded | NA | NA | Bomb fishing,<br>coral mining | Hard coral<br>cover = 23% | 25 |
| Indonesia | D4 | Degraded | NA | NA | Bomb fishing,<br>coral mining | Hard coral<br>cover = 2% | 25 |
| Indonesia | D5 | Degraded | NA | NA | Bomb fishing,<br>coral mining | Hard coral<br>cover = 3% | 28 |
| Indonesia | D6 | Degraded | NA | NA | Bomb fishing,<br>coral mining | Hard coral<br>cover = 16% | 24 |
| Indonesia | H1 | Healthy | NA | NA | NA | Hard coral<br>cover = 74% | 26 |
| Indonesia | H2 | Healthy | NA | NA | NA | Hard coral<br>cover = 87% | 25 |
| Indonesia | H3 | Healthy | NA | NA | NA | Hard coral<br>cover = 71% | 27 |
| Indonesia | H4 | Healthy | NA | NA | NA | Hard coral<br>cover = 57% | 5 |
| Indonesia | H5 | Healthy | NA | NA | NA | Hard coral<br>cover = 75% | 6 |
| Indonesia | H6 | Healthy | NA | NA | NA | Hard coral<br>cover = 67% | 25 |
| Indonesia | R1 | Mid-stage<br>restored | Approx.<br>600 | 52 months | Bomb fishing,<br>coral mining | Hard coral<br>cover = 83% | 25 |
| Indonesia | R2 | Mid-stage<br>restored | Approx.<br>600 | 48 months | Bomb fishing,<br>coral mining | Hard coral<br>cover = 78% | 24 |
| Indonesia | R3 | Mid-stage<br>restored | Approx.<br>600 | 41 months | Bomb fishing,<br>coral mining | Hard coral<br>cover = 53% | 25 |
| Indonesia | R4 | Mid-stage<br>restored | Approx.<br>600 | 36 months | Bomb fishing,<br>coral mining | Hard coral<br>cover = 72% | 26 |
| Indonesia | R5 | Mid-stage restored | Approx. 600 | 33 months | Bomb fishing, coral mining | Hard coral cover = 48% | 26 |
| Indonesia | R6 | Mid-stage restored | Approx. 600 | 53 months | Bomb fishing, coral mining | Hard coral cover = 58% | 26 |
| Indonesia | N1 | Early-stage restored | Approx. 600 | 1 months | Bomb fishing, coral mining | Approx. 15% | 6 |
| Indonesia | N2 | Early-stage restored | Approx. 600 | 3 months | Bomb fishing, coral mining | Approx. 15% | 6 |
| Indonesia | N3 | Early-stage restored | Approx. 600 | 1 months | Bomb fishing, coral mining | Approx. 15% | 7 |
| Kenya | D1 | Degraded | NA | NA | Overfishing | Hard coral cover = 0% | 30 |
| Kenya | D2 | Degraded | NA | NA | Overfishing | Hard coral cover = 1% | 35 |
| Kenya | H1 | Healthy | NA | NA | NA | Hard coral cover 54% | 4 |
| Kenya | H2 | Healthy | NA | NA | NA | Hard coral cover = 54% | 35 |
| Kenya | N1 | Early-stage restored | 120 | 3 months | Overfishing | Hard coral cover = 17% | 35 |
| Maldives | D1 | Degraded | NA | NA | Construction sedimentation | Fish biomass = 21kg/ha<sup>-1</sup>, spp. richness = 15.5 | 34 |
| Maldives | D2 | Degraded | NA | NA | Construction sedimentation | Fish biomass = 46kg/ha<sup>-1</sup>, spp. richness = 15.5 | 34 |
| Maldives | H1 | Healthy | NA | NA | NA | Fish biomass = 94kg/ha<sup>-1</sup>, spp. richness = 31 | 35 |
| Maldives | H2 | Healthy | NA | NA | NA | Fish biomass =<br>124kg/ha<sup>-1</sup>, spp.<br>richness = 29 | 35 |
| Maldives | N1 | Early-stage<br>restored | 101 | 2 months | Construction<br>sedimentation | Fish biomass =<br>54kg/ha<sup>-1</sup>, spp.<br>richness = 19.5 | 41 |
| Mexico | D1 | Degraded | NA | NA | Overfishing,<br>pollution,<br>disease | Hard coral<br>cover < 5% | 32 |
| Mexico | D2 | Degraded | NA | NA | Overfishing,<br>pollution,<br>disease | Hard coral<br>cover < 5% | 32 |
| Mexico | H1 | Healthy | NA | NA | NA | Hard coral<br>cover = 29% | 32 |
| Mexico | H2 | Healthy | NA | NA | NA | Hard coral<br>cover = 14% | 29 |
| Mexico | H3 | Healthy | NA | NA | NA | Hard coral<br>cover = 17% | 31 |
| Mexico | R2 | Mid-stage<br>restored | 350 | 35 months | Overfishing,<br>pollution,<br>disease | Hard coral<br>cover = 25% | 30 |
| Mexico | N1 | Early-stage<br>restored | 139 | 3 months | Overfishing,<br>pollution,<br>disease | Hard coral<br>cover = 14% | 32 |

**Supplementary Table 2:**
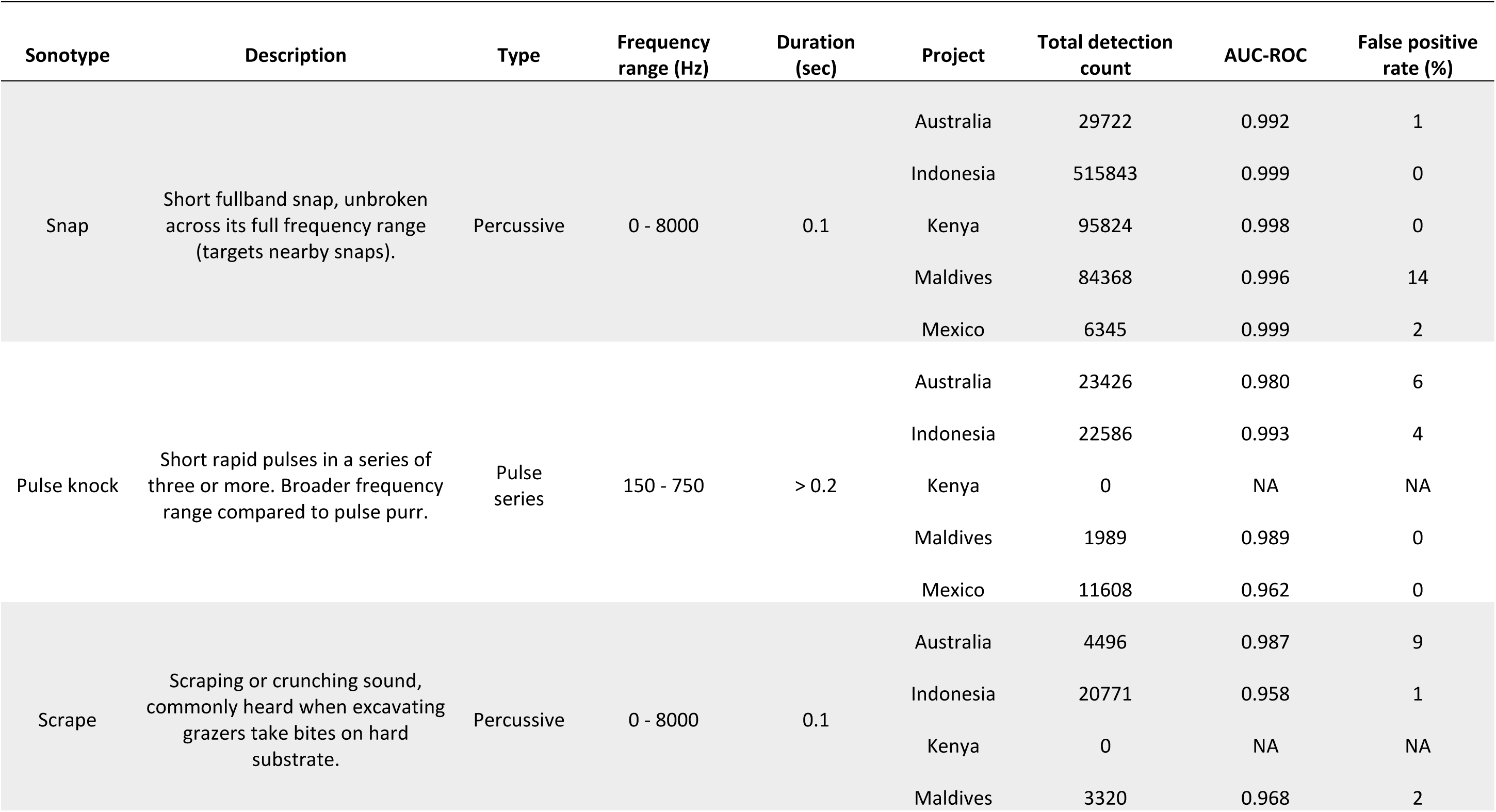
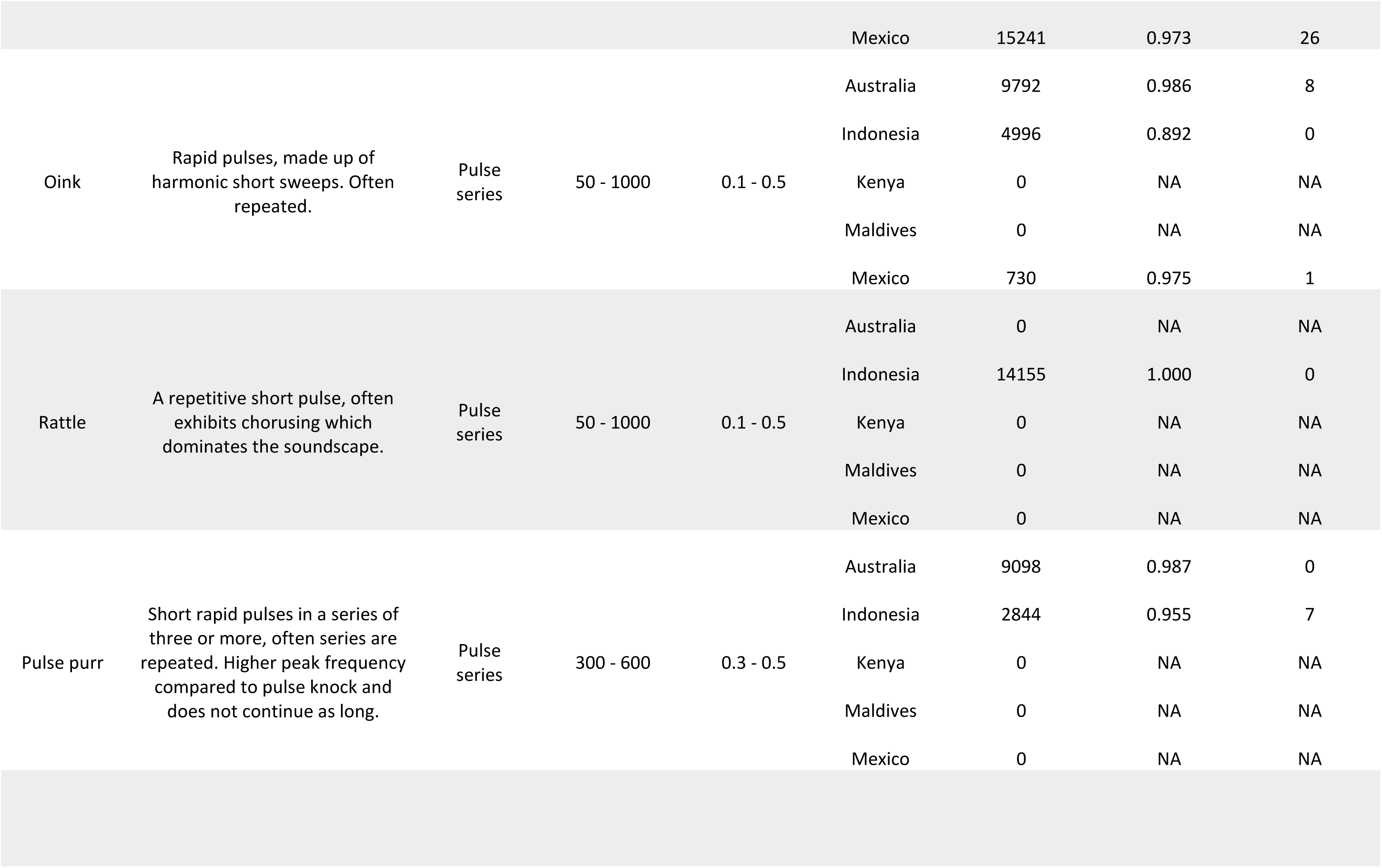
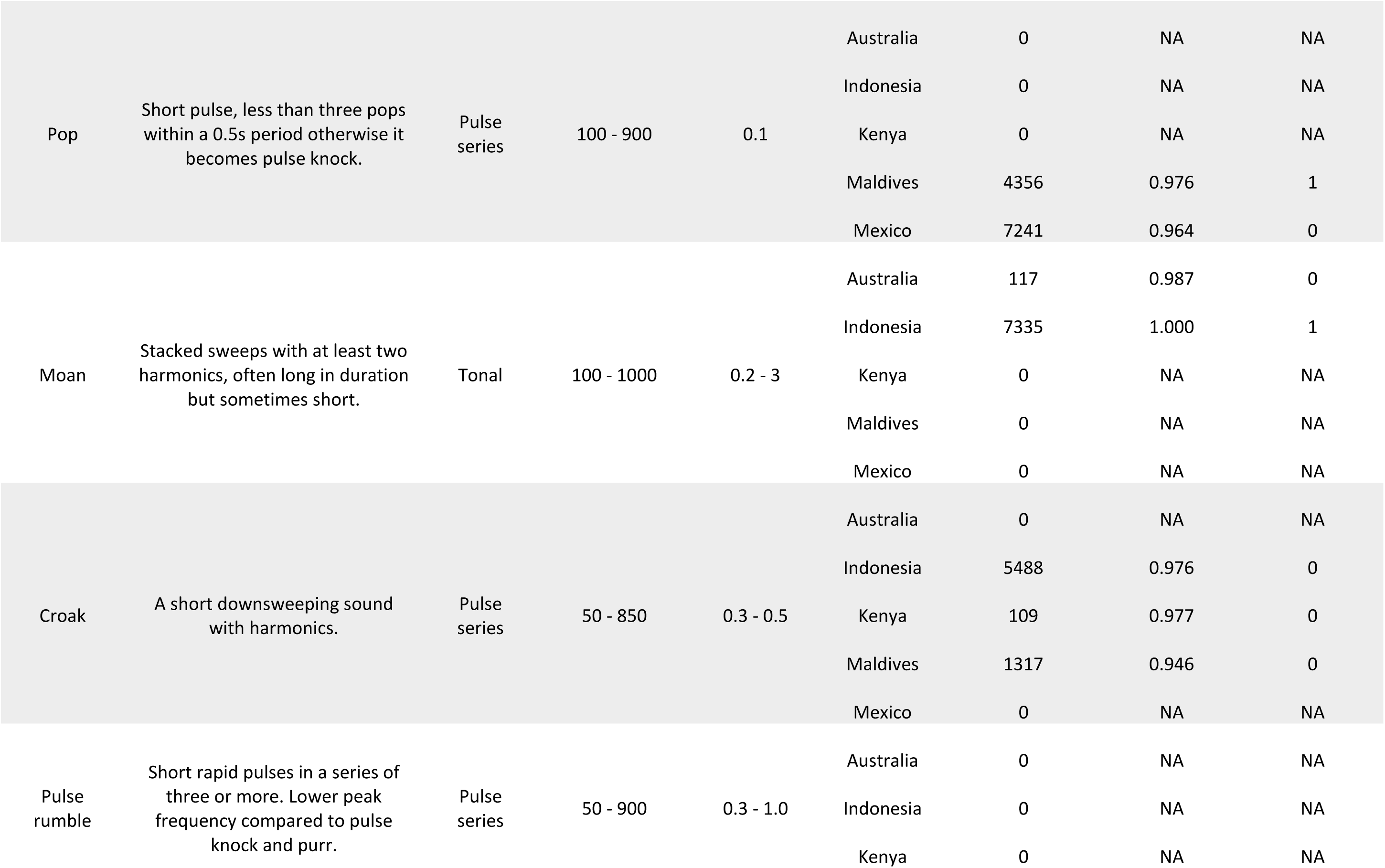
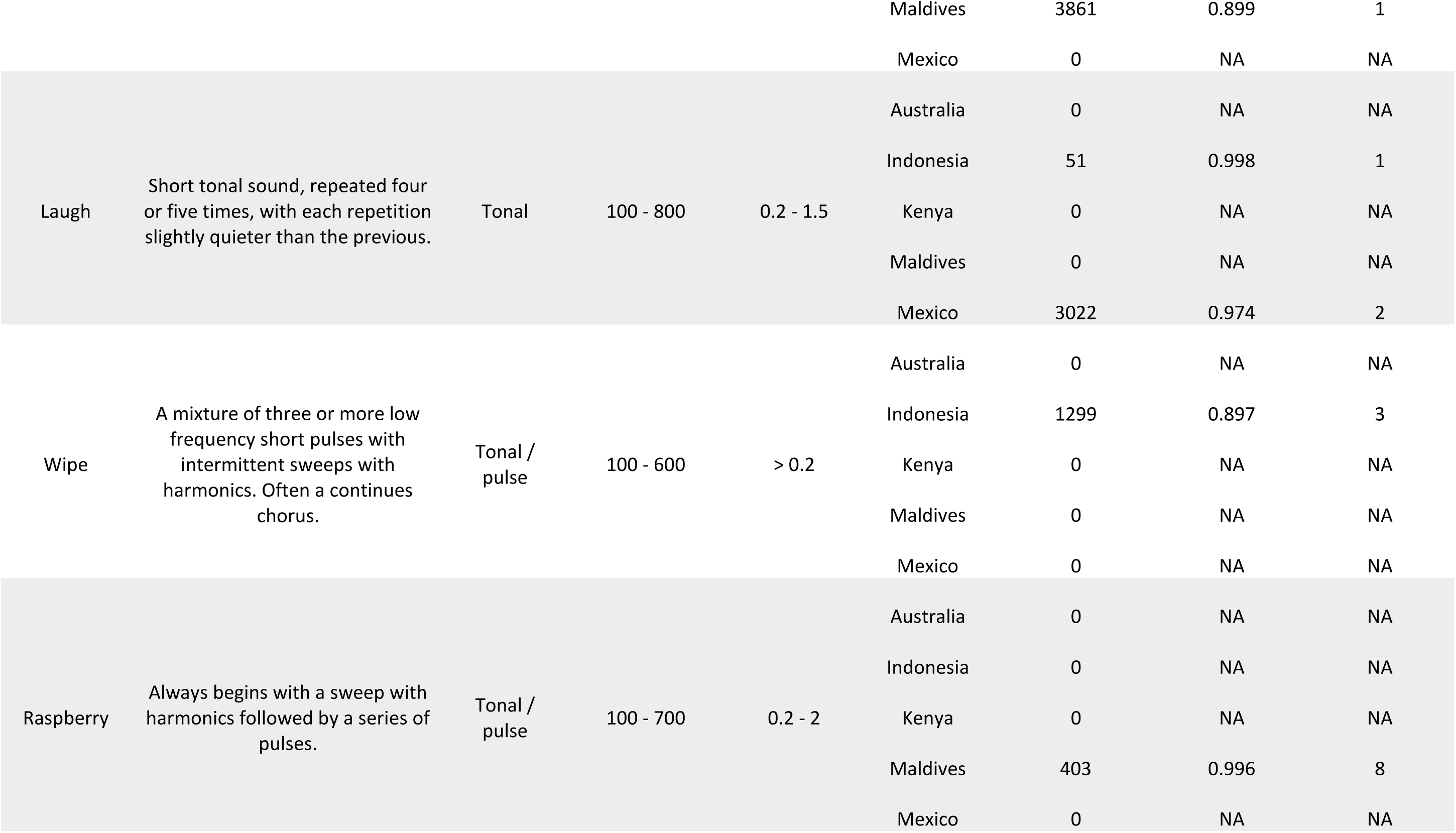
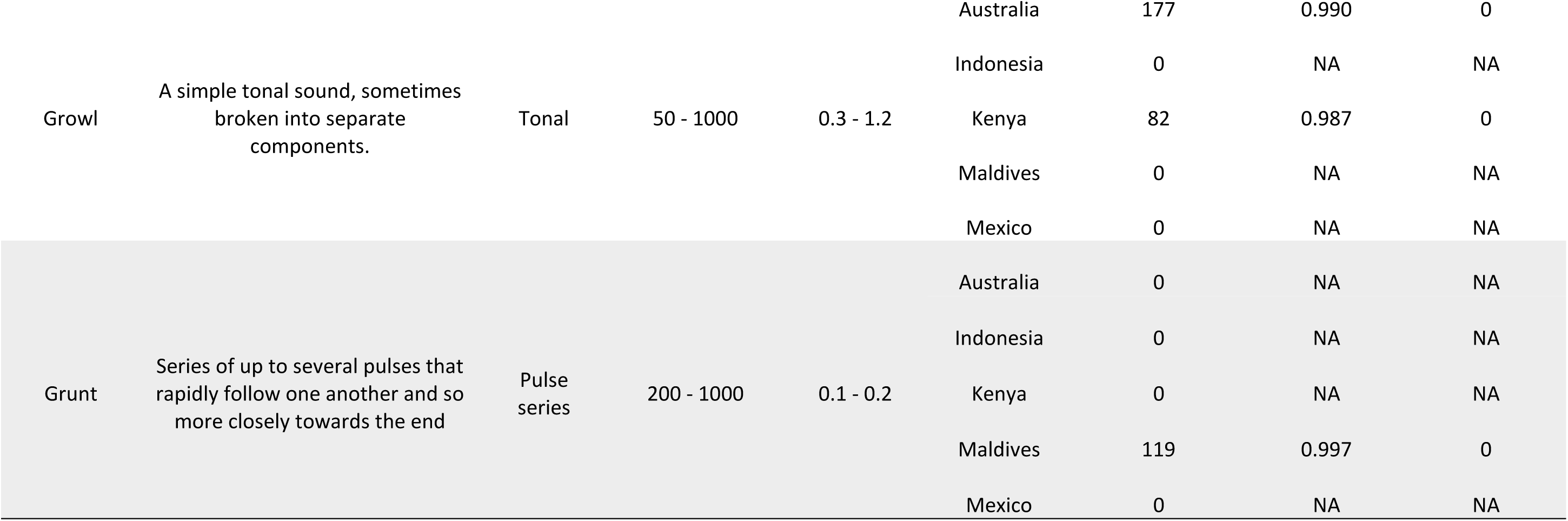
Descriptions and characteristics of all sonotypes identified during the study. Frequency range (Hz) and duration (sec) represent the minimum and maximum bounds of each. Where total detection count is 0, the sonotype was not found to be present within the project during the initial screening and therefore no classifier was trained to detect this.

